# Ecological niche adaptation of a bacterial pathogen associated with reduced zoonotic potential

**DOI:** 10.1101/2020.09.09.288845

**Authors:** Mark Kirkwood, Prerna Vohra, Matt Bawn, Gaëtan Thilliez, Hannah Pye, Jennifer Tanner, Cosmin Chintoan-Uta, Priscilla Branchu, Liljana Petrovska, Timothy Dallman, Neil Hall, Mark P. Stevens, Robert A. Kingsley

## Abstract

The emergence of new bacterial pathogens is a continuing challenge for agriculture and food safety. *Salmonella enterica* serovar Typhimurium (*S*. Typhimurium) is a major cause of foodborne illness worldwide, with pigs a major zoonotic reservoir. Two variants, *S*. Typhimurium phage type U288 and monophasic *S*. Typhimurium (*S*. 4,[5],12:i:-) ST34 emerged and have accounted for the majority of isolates from pigs in the UK in the past two decades, but have distinct host range and risk to food safety. ST34 accounts for over 50% of all *S*. Typhimurium infections in people while U288 less than 2%. U288 and ST34 form distinct phylogenetic clusters within *S*. Typhimurium, defined by approximately 600 SNPs within their 5 Mbp genomes. Evolution of the U288 clade from an LT2-like ancestor was characterised by the acquisition of AMR genes, insertions and deletions in the virulence plasmid pU288-1, and the accumulation of polymorphisms, some of which resulted in truncation of coding sequences. U288 isolates exhibited lower growth rate and viability following desiccation compared to ST34 isolates, characteristics that could affect transmission through the food chain. U288 and ST34 isolates exhibited distinct outcomes of infection in the murine model of colitis, and colonised pigs in a manner that affected the disease symptoms and distribution in organs. U288 infection was more disseminated in the lymph nodes while ST34 were recovered in greater numbers in the intestinal contents. These data are consistent with the evolution of *S*. Typhimurium U288 adaptation to pigs that may determine their reduced zoonotic potential.

**Importance:** Bacterial pathogens continually evolve to exploit new ecological niches as they emerge due to human activity including agricultural, medical or societal practice. The consequences of the emergence of new pathogens may affect outcome of infection and risk to human or animal health. Genome sequence can resolve the population structure, identify variants that are evolving as they enter a new niche, and pinpoint potential functional divergence. We report a variant *S*. Typhimurium that adapted to a unique niche distinct to that occupied by a second *S*. Typhimurium variant circulating in the same pig populations. Adaptation was accompanied by phenotypic and genotypic changes consistent with a more invasive lifestyle and a decreased zoonotic potential observed in the epidemiological record. Our findings suggest that pathogen genotypic variation may be useful in estimating zoonotic potential and threat to livestock welfare.

## Introduction

Emergence of infectious diseases presents new challenges for the management of human and livestock health, with substantial human and economic costs through excess mortality and morbidity, and lost productivity. The emergence of 335 human infectious diseases between 1945 and 2004 was dominated by zoonoses and bacterial aetiological agents, in-part due to the emergence of new variants of drug resistant pathogens (1). Emergent niches are rapidly occupied by bacteria from the vast genetic diversity in the biosphere, that are best suited to its exploitation. The plastic genomes, due to horizontal gene transfer and errors during replication, and relatively short replication time, facilitate rapid evolution by the selection of variants with increased survival and replication (2).

The genus *Salmonella* consists of over 2500 different serovars, such as Typhimurium, Enteritidis and Choleraesuis, that have diverse host ranges, pathogenicity and risk to human health. Prevalent serovars differ between host species and in different geographical locations. The prevalence of each serovar may also vary over time, although certain serovars are often characteristic of specific host species and maintained for decades. For example, *S*. Typhimurium (including monophasic variants) has consistently been a dominant serovar in pigs globally and accounts for around two thirds of isolates in the UK (3). Despite the apparently stable prevalence of *S*. Typhimurium in pig populations over decades, the epidemiological record of variants identified by phage type, indicates a dynamic process where distinct phage types increase and decrease in prevalence over time (3). The epidemiology of *S*. Typhimurium in human infections in Europe since the middle of the 20^th^ century was characterized by successive emergence and replacement of dominant phage types including DT9, DT204, DT104 and most recently monophasic *S*. Typhimurium (*S*. 1,4,[5],12:i:-) with sequence type 34 (ST34) that is predominantly DT193/DT120. At their peak incidence, each accounted for over half of all human isolates of *S*. Typhimurium. In UK pigs, since around the year 2003, isolates of U288 and DT193 have dominated (3). U288 appeared in UK pig populations around 2003 followed around the year 2006 by monophasic *S*. Typhimurium (*S*. 1,4,[5],12:i:-) with sequence type (ST34) rapidly emerging in pig populations around the world (3, 4). These two variants co-existed in the pig population and together accounted for around 80% of isolates (5). Despite, approximately half of all pork consumed in the UK being from UK pig herds (6), throughout this period U288 were rarely isolated from human infections (7). In contrast, monophasic *S*. Typhimurium ST34 isolations from human infections closely reflected its prevalence in pig populations and by the year 2013, over half of all *S*. Typhimurium isolates from human non-typhoidal salmonellosis (NTS) in the UK were monophasic *S*. Typhimurium ST34 (8, 9). A similar epidemiology has been reported in other European countries (10), where the remaining pork consumed in the UK is sourced.

Non-typhoidal Salmonella (NTS) causes up to 78 million human infections each year, globally (11) and pigs are a major zoonotic reservoir, with 10-20% of human salmonellosis in Europe attributable to pigs (12, 13). A baseline survey reported prevalence of 21.2% and 30.5% in mesenteric lymph nodes and caecal contents, respectively, for UK slaughter pigs (14, 15). It is believed that contamination of pig carcasses with faeces and gut contents at slaughter, and the ability of *Salmonella* to spread from the gut to other organs, results in contamination of meat products that enter the food chain and pose a risk to humans if improperly handled or cooked. However, the relative risk from contamination of meat by gut contents during slaughter or from tissue colonised by *Salmonella* prior to slaughter is not known. Survival in food depends upon adaptive response to environmental stresses including osmotic pressure from preservatives and desiccation, antimicrobial activity of preservatives and fluctuating temperatures during storage or cooking. In order to cause disease, *Salmonella* may also need to replicate to achieve a population size able to overcome the colonisation resistance of the host.

Multiple pathovariants of *S*. Typhimurium evolved from a broad host range ancestor resulting in distinct host range, outcome of infection and risk to food safety (16–18). An understanding of the molecular basis of risk to food safety of pathovariants is critical to improve assessment of risk and devise intervention strategies aimed at decreasing the likelihood that *Salmonella* is present in food intended for human consumption. Moreover, analysis of *Salmonella* genome sequences by machine learning has enabled the prediction of host of origin for specific variants, which may aid source attribution in outbreak investigations (18–20). We therefore investigated the population structure of *S*. Typhimurium U288 and the genomic evolution leading to the clonal expansion of U288 associated with porcine infection by analysis of whole genome sequences. We compared the interaction of representative strains of the *S*. Typhimurium U288 and the monophasic *S*. Typhimurium epidemic clades with the environment and *in vivo* in the pig host to gain insight into the phenotypic consequences of their distinct evolutionary trajectories.

## Materials and Methods

### Bacterial strains and culture

*Salmonella* Typhimurium U288 and ST34 isolates used in this study were isolated from human clinical infections during routine diagnostic testing by Public Health England (PHE), or from animals during routine surveillance or epidemiological investigation by Animal and Plant Health Agency (APHA) (Supplementary Table 1). All sequence data generated in this study is available in SRA database under BioProject accession number PRJNA641292. 131 genotypically diverse *S*. Typhimurium isolates used to place U288 and ST34 in phylogenetic context have been described previously (21, 22). Characteristics and metadata of isolates are summarised in Supplementary Table 1. Bacterial isolates were stored at −80°C in 25% glycerol and routinely cultured overnight in 5 mL LB broth at 37°C with shaking at 200 rpm, or on solid medium consisting of Luria Bertani (LB) broth or MacConkey containing 5% Agar, and supplemented with chloramphenicol (30 mg/l) or kanamycin (50 mg/l) as appropriate.

### Construction of wild type isogenic tagged strains and single knock out

A modified recombineering method based on the Lambda Red system was used to construct knockout mutations in *S.* Typhimurium SL1344 and wild type isogenic tagged strains (WITS) (23). Briefly, primers where used to amplify the kanamycin resistance cassette from pKD4 and recombination into the genome was directed by the inclusion of 50 nucleotide sequence flanking the insertion site (Supplementary Table 2). For the construction of WITS, insertion was directed to the intergenic region of *iciA* and *yggE* at position 3,247,245 in SL1344 (24). Each forward primer included a unique 10 nucleotide tag which allowed identification of the tagged strain through whole genome sequencing.

### Determination of growth rate, biofilm formation and desiccation survival

For determination of growth rate, bacterial cultures in LB broth were diluted to approximately 1×10^5^ CFU per ml and incubated at 37°C with atmospheric aeration or an anaerobic environment (10% CO_2_, 5% H_2_, %% O_2_ and 80% N_2_) and viable bacteria in colony-forming units (CFU) enumerated by serial dilution and culture on LB agar at 1, 3, 5 and 7 hours post inoculation. Doubling time was calculated in the exponential range of growth using the mean from three biological replicates. Determination of survival after desiccation was based on a method previously described (25). Briefly, bacteria were cultured in LB broth, harvested by centrifugation, washed with phosphate-buffered saline pH7.4 (PBS) and re-suspended in PBS and adjusted to OD_600nm_ of 1. Next, 0.05 ml of cell suspension were added to polystyrene 96 well plate (Nunc) and desiccated at 22°C, 36% relative humidity (RH). Desiccated plates were stored in a sealed vessel containing saturated potassium acetate solution, to maintain RH at 36% and incubated at 22°C for 24 hours. Cells were re-suspended in 0.2 ml PBS and viable counts were determined by culture of serial 10-fold dilutions on LB agar. Colony counts were determined at 0 hours and 24 hours and a percentage survival calculated from at least three biological replicates. To study biofilm formation by bacterial isolates, liquid cultures were incubated statically in polystyrene 96-well plates at 22°C for 24 hours, washed once with PBS pH7.4, and attached bacteria were stained with crystal violet and absorbance read at 340nm. Data points represent the mean of at least three biological replicates.

### Preparation of genomic DNA and sequencing

Genomic DNA for short-read sequencing was extracted using Wizard® Genomic DNA Purification (Promega) from a culture inoculated form a single colony and incubated for 18 hours at 37°C. Low Input, Transposase Enabled (LITE) Illumina compatible libraries were constructed using a modified protocol based on the Illumina Nextera kit (Illumina, California USA). A total of 1ng of DNA was combined with 0.9 μl of Nextera reaction buffer and 0.1 μl Nextera enzyme in a reaction volume of 5 μl and incubated for 10 minutes at 55°C. Following the initial incubation, 2.5 μl of 2 μM custom barcoded P5 and P7 compatible primers, 5 μl 5x Kapa Robust 2G reaction buffer, 0.5 μl 10mM dNTPs, 0.1 μl Kapa Robust 2G enzyme and 10.4 μl water were mixed and amplified by incubating the sample for 72°C for 3 minutes, followed by 14 PCR cycles consisting of 95°C for 1 minute, 65 °C for 20 seconds and 72 °C for 3 minutes. 20μl of amplified DNA was added to 20μl of Kapa beads and incubated at room temperature for 5 minutes to precipitate DNA molecules >200bp onto the beads. The beads were then pelleted on a magnetic particle concentrator (MPC), the supernatant removed, and two 70% ethanol washes performed. After removal of the final 70% ethanol wash, the beads were left to dry for 5 minutes at room temperature before the beads were re-suspended with 20μl of 10mM Tris-HCl, pH8. This was then incubated at room temperature for 5 minutes to elute the DNA molecules. The tube was then placed back on the MPC, the beads allowed to pellet, and the aqueous phase containing the size selected DNA molecules transferred to a new tube. The size distribution of each purified library was determined on a PerkinElmer GX by diluting 3 μl in 18 μl 10mM Tris-HCl, pH8. Using the PerkinElmer GX software, the smear analysis function was used to determine the amount of material in the 400 to 600bp size range and this information used to equimolar pool purified libraries. Once pooled the samples were then subjected to size selection on a Sage Science 1.5% BluePippin cassette recovering molecules between 400 and 600bp. QC of the size selected pool was performed by running 1μl aliquots on a Life Technologies Qubit high sensitivity assay and an Agilent DNA High Sense BioAnalyser chip and the concentration of viable library molecules measured using qPCR. 10pM library pools were loaded on a HiSeq4000 (Illumina, California, USA) based on an average of the qubit and qPCR concentrations using a mean molecule size of 425bp.

### Phylogenetic reconstruction and time-scaled inference

Paired-end sequence files for each isolate were mapped to the SL1344 reference genome (FQ312003) (24) or S01906-05 (PRJEB34597) (18) using the Rapid haploid variant calling and core SNP phylogeny pipeline SNIPPY (version 3.0) (https://github.com/tseemann/snippy). The size of the core genome was determined using snp-sites (version 2.3.3) (26), outputting monomorphic as well as variant sites and only sites containing A,C,T or G. Variant sites were identified and a core genome variation multifasta alignment generated. The sequence alignment of variant sites was used to generate a maximum likelihood phylogenetic tree with RAxML using the GTRCAT model implemented with an extended majority-rule consensus tree criterion (27). The genome sequence of *S.* Heidelberg (NC_011083.1) was used as an outgroup in the analysis to identify the root and common ancestor of all *S*. Typhimurium strains.

To infer the time of nodes on phylogeny, we used the BactDating software package implemented in R (28), in which sequence variation in the core genome was analysed with potential regions of recombination removed using Gubbins (29). The resulting sequence alignments were used to construct a maximum likelihood phylogenetic tree using RAxML and the true root was estimated using *S*. Typhimurium SL1344 genome as an outgroup. The MCMC was run for 1 million iterations and the convergence and mixing of chains were 113.3, 129.5, 145.8 for (□, □, □, respectively) calculated using the R package corda (30).

### Pan-genome analysis and *In-silico* genotyping

The pan-genome of U288, ST34 and representative strains of S. Typhimurium was determined as described previously (18). The presence of antibiotic resistance, virulence and plasmid replicon genes in short-read data was determined by the mapping and local assembly of short reads to data-bases of candidate genes using Ariba (31). The presence of candidate genes from the ResFinder (32), VFDB (33) and PlasmidFinder (34) databases was determined. Reads were mapped to candidate genes using nucmer with a 90% minimum alignment identity. This tool was also used to determine the presence of specific genes or gene allelic variants. The results of the ARIBA determination of the presence or absence of the *lpfD* gene were confirmed using SRST2 (35) setting each alternative form of the gene as a potential allele. SRST2 was also used to verify the ARIBA findings of the VFDB data set, as the presence of orthologous genes in the genome was found to confound the interpretation of results.

### Metabolic profiling using OMNIlog microarray system

To assess utilisation of carbon sources, isolates were cultured on LB agar overnight at 37°C, inoculated into IF-0 medium containing tetrazolium dye, and added to a PM-1 plate (with 95 different carbon sources), according to the manufacturer’s instructions (IBiolC). Accumulation of purple indicator dye as a consequence of redox activity was measured every 15 minutes for 48 hours. Raw absorbance data were processed using R software and the opm package (36). The area-under-the-curve was a metric for total respiration of the indicated carbon sources and plotted as a heatmap.

### Epithelial cell invasion assays

IPEC-J2 porcine epithelial cells and T84 human epithelial cells were cultured and routinely passaged in Dulbecco Modified Eagles Medium (DMEM), containing high glucose or low glucose for each cell line, respectively. 24-well tissue culture plates were seeded with 1 × 10^5^ cells/ml of epithelial cell lines, and incubated at 37°C 5% CO_2_ overnight. Stationary phase cultures of *S.* Typhimurium were balanced to OD_600nm_ ~1.0 in PBS and used to inoculate epithelial cells at a multiplicity of infection (MOI) of 20. Cells were incubated for 30 minutes before washing 5 times with PBS and re-suspending in DMEM + gentamicin (100 mg/l) and incubating for a further 30 minutes at 37°C in 5% CO_2_ atmosphere to kill extracellular bacteria. The medium was then replaced with DMEM + gentamicin (10 mg/l) and the plates were incubated for 60 minutes at 37°C in 5% CO_2_ atmosphere. Plates were then incubated at 37°C in 5% CO_2_ atmosphere for 90 minutes, before a final wash with PBS + 0.1% Triton-X100. Plates were left for 2 minutes and cells were disrupted by vigorous pipetting for one minute followed by serial dilution and culture on LB agar to determine viable counts.

### Streptomycin pre-treated mouse infections

Groups of five female 6-9 week old C57bl/6 mice were treated with 20 mg of streptomycin sulphate by oral gavage 24 hours prior to inoculation of *S*. Typhimurium. *S*. Typhimurium strains were cultured for 18 hours in 50ml LB broth and approximately 5×10^6^ to 1×10^7^ CFU were inoculated orally in 0.2 ml of PBS pH7.4 by gavage. On day 3 post-inoculation mice were humanly euthanised and the cecum was aseptically removed. An approximately 3mm section of the cecum half way down the organ from the ileal junction were removed and fixed in formalin for histopathology examination of 5um thin sections stained with hemotoxylin and eosin. Approximately 5 mg of cecum tissue was removed and placed in RNAlater and stored at −80°C (Thermo Fisher). The remaining cecum (approximately two thirds) was homogenised in sterile PBS pH7.4 and serial dilutions plated on LB agar containing 0.05 mg/ml kanamycin, incubated at 37°C for 18 hours and viable counts enumerated.

### Determination of Nos2 and Cxl1 expression in mouse cecum tissue

RNA was prepared using Guanidinium thiocyanate-phenol-chloroform extraction with Tri Reagent (Merck). Tissue was homogenized in 1 ml of Tri reagent and disrupted using 1.4mm zinc oxide beads in a bead beater and tissue debris removed by centrifugation. 0.2 ml of nuclease free water and 0.2ml of chloroform were added to the supernatant, mixed and centrifuged for 15 minutes at 12,000 × g. 0.5 ml of isopropanol was added to the upper aqueous phase and centrifuged for 15 minutes at 12,000 × g. The resulting RNA pellet was washed twice with 70% ethanol, briefly dried and resuspended in 0.02 ml of RNase free water. The relative abundance of Nos2 and Cxl1 mRNA was determined by quantitative RT-PCR using primers specific to the test genes and the Gapdh house-keeping gene as the control (Supplementary Table ?) as described previously (37).

### Mixed-strain infection of pigs

Four 6-week-old Landrace x Large White x Durock pigs were challenged orally with a mixed-strain inoculum as previously described (38). Animal experiments were conducted according to the requirements of the Animals (Scientific Procedures) Act 1986 (project licence PCD70CB48) with the approval of the local ethical review committee. Pigs were confirmed to be culture negative for *Salmonella* before inoculation by enrichment of faecal samples in Rappaport-Vassiliadis broth at 37°C for 18 hours, followed by plating on MacConkey agar at 37°C for 24 hours. A mixed-strain inoculum was prepared by mixing equal volumes of individual cultures of the six strains grown statically at 37°C for 16 hours in LB broth supplemented with 50 mg/l kanamycin, which were optical density (OD_600nm_) standardized to contain 8.9 log_10_ CFU/ml. This was confirmed retrospectively by plating 10-fold serial dilutions on MacConkey agar. Aliquots of the inoculum were stored at −20°C for DNA extraction. Five ml of the mixed-strain inoculum was mixed with 5 ml of antacid [5% Mg(SiO_3_)_3_, 5% NaHCO_3_, and 5% MgO in sterile distilled water] to promote colonization and administered orally by syringe before the morning feed. Pigs were fed as normal following challenge. Rectal temperatures were recorded every 24 hours and faecal samples were collected at 24 and 48 hours post-infection. At 72 hours post-infection, a section of distal ileal mucosa, mesenteric lymph nodes (MLNs) draining the distal ileal loop, a section of spiral colon, colonic lymph nodes (CLNs) and a section of liver were collected. Lymph nodes were trimmed of excess fat and fascia, and the sections of distal ileum and spiral colon were washed gently in PBS to remove nonadherent bacteria. One gram of each tissue was homogenized in 9 ml of PBS in gentleMACS M tubes using the appropriate setting on the gentleMACS dissociator (Miltenyi Biotec). Homogenates were filtered through 40-μm-pore-size filters and an aliquot was used to determine viable counts. The remaining homogenate was spread onto 10 MacConkey agar plates (500 μl per plate) and incubated overnight at 37°C. The bacterial lawns recovered from each sample were collected by washing with PBS, and the pellets were stored at −20°C for DNA extraction. Genomic DNA (gDNA) was extracted from the pellets using the NucleoSpin tissue kit (Macherey-Nagel), according to the manufacturer’s instructions. The quality and quantity of DNA were assessed initially by NanoDrop 3300 (Thermo Scientific), and samples with an *A*_260/280_ of <=1.8 were considered suitable for library preparation. These were confirmed further by using the DNA ScreenTape (Agilent Technologies) and the Qubit double-stranded DNA (dsDNA) BR assay kit (Life Technologies), respectively. One microgram of gDNA with a DNA integrity number (DIN) of <=6 was used for library preparation using the TruSeq PCR-free library preparation kit (Illumina) according to the manufacturer’s protocol. Whole genome sequencing on the HiSeq system (Illumina) followed by bioinformatics analysis were performed as previously described (38), with the exception that strains were quantified by mapping sequence data to the unique WITS tag sequence incorporated chromosomally into each strain. Sequence data was submitted to the NCBI SRA database (Supplementary Table 3). For each strain, the percentage in a population was calculated as the average WITS frequency x 100. Data are presented as the mean ± standard error of the mean (SEM). Raw sequence data were deposited in the Biosample database at NCBI (Supplementary Table 3).

### Single-strain infection of pigs

From the strain phenotypes identified in the mixed-strain infection, one representative strain of each DT193 and U288 was selected for *in vivo* phenotype validation, S04698-09 and 11020-1996, respectively. The strains were grown statically at 37°C for 16 hours in LB broth supplemented with 50 mg/l kanamycin and the optical densities (OD_600nm_) were standardized to contain 9.2 log_10_ CFU/ml, which was confirmed retrospectively by plating 10-fold serial dilutions on MacConkey agar. Groups of four *Salmonella*-free pigs were challenged orally with 5 ml of each strain as described above. Rectal temperatures were recorded every 12 hours and faecal samples were collected every 24 hours post-infection. Clinical scores were calculated for each animal using their temperatures, physiological signs and faecal consistency. At 72 hours post-infection, tissue samples were collected and processed for viable counts as described above. The bacterial load of each strain in each tissue of the infected pigs was determined. Data are presented as the mean ± SEM.

### Statistical analysis

Statistical tests were performed in GraphPad Prism version 8.00 (GraphPad Software). The viable counts of bacteria in each case is presented as mean ± SEM, and differences between strains were analysed using the Mann-Whitney test of significance. Area under the curve analysis followed by the Mann-Whitney test was used to analyse the cumulative clinical scores of the infected pigs during single-strain infections. *P* values of ≤0.05 were considered to be statistically significant.

## Results

### *S*. Typhimurium U288 and monophasic *S*. Typhimurium ST34 exhibit distinct host range

The epidemiological record indicates that *S*. Typhimurium U288 was first reported in pigs in the UK around the year 2000 and thereafter became the dominant phage type isolated for much of the following decade (3). Monophasic *S*. Typhimurium ST34 emerged around seven years later in UK pigs, and these two variants have co-existed in pig populations since. Retrospective analysis of the frequency of U288 and monophasic *S*. Typhimurium ST34 isolated from animals in the UK by the Animal and Plant Health Agency (APHA) between 2006 and 2016, revealed distinct host ranges (Figure 1). During this period, a total of 1535 and 2315 isolates of *S*. Typhimurium U288 and monophasic *S*. Typhimurium, were reported by APHA from animals in the UK, respectively. *S*. Typhimurium U288 was almost exclusively isolated from pigs, while in contrast, monophasic *S*. Typhimurium, although predominantly isolated from pigs (39), was also isolated from multiple host species including cattle and poultry populations (Figure 1).

**Figure 1.**
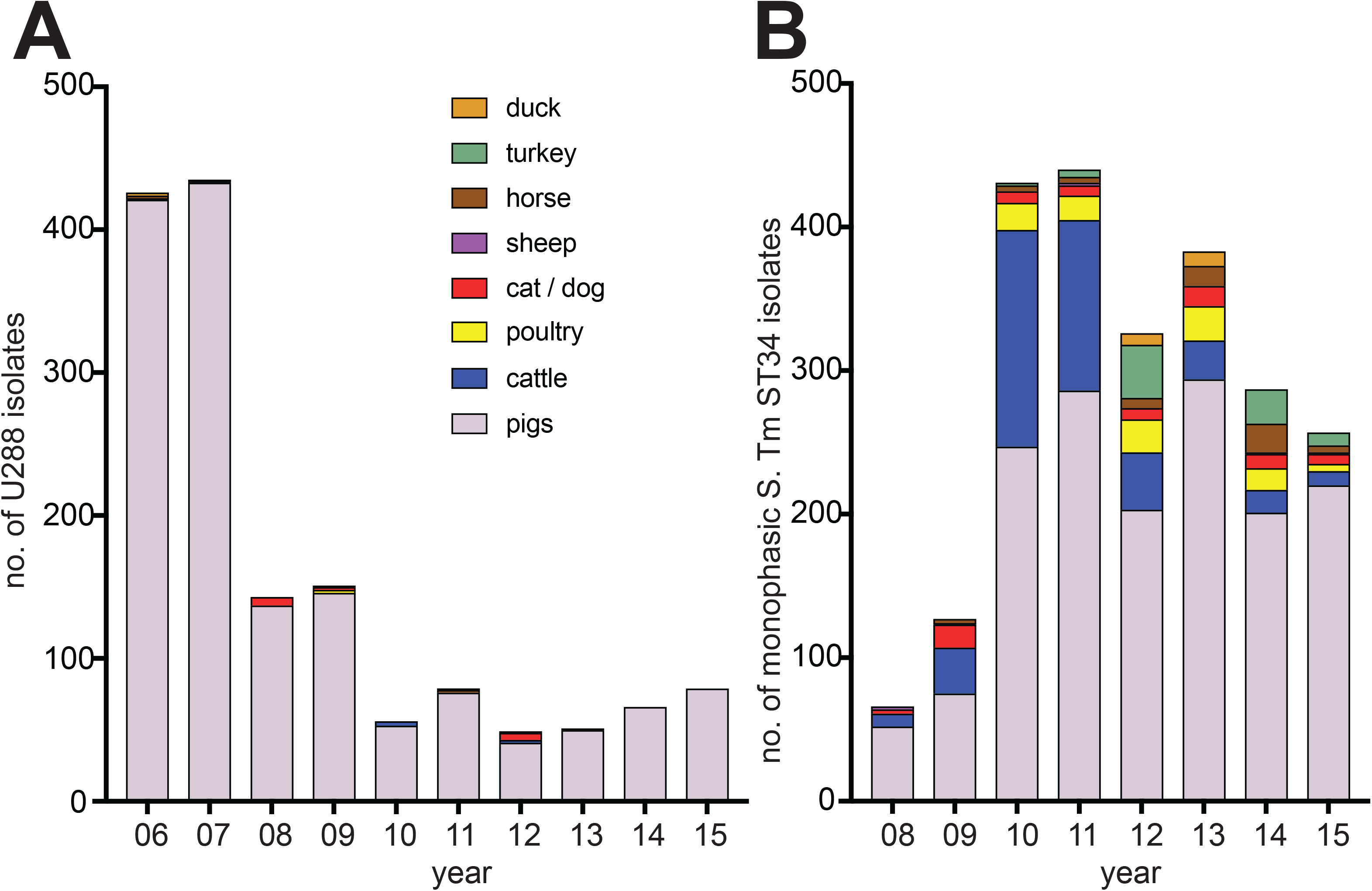
Animal species source of monophasic *S*. Typhimurium ST34 and *S*. Typhimurium U288. Stacked bar chart indicating the animal source (see colour key inset) of *S*. Typhimurium U288 (A) and monophasic S. Typhimurium ST34 (B) isolated in England and Wales by Animal and Plant Health Agency 2006-2015.

### The *S*. Typhimurium U288 and ST34 isolates form distinct phylogroups

To investigate the phylogenetic relationship of *S*. Typhimurium U288 isolates, we first constructed a maximum likelihood tree using variation in the recombination-purged core-genome sequence of 1666 *S*. Typhimurium isolates from human clinical infections in England and Wales between April 2014 and December 2015 for which both whole genome sequence and the phage type data were available; 24 (1.4%) were U288. Of these, 20 of 24 *S*. Typhimurium U288 isolates were present in a distinct clonal group, composed of 33 isolates in total (henceforth referred to as the U288 clade, Supplementary Figure 1). The remaining 13 isolates within the predominantly U288 clade were isolates of phage types DT193 (5 isolates), U311 (3 isolates), U302 (1 isolate), or reacted did not conform (RDNC, 4 isolates). The main U288 clade was closely related to 13 human clinical isolates, of various phage types, but predominantly U311; none were U288.

To investigate the relationship of contemporaneous *S*. Typhimurium U288 in the UK pig population and human clinical isolates, we determined the whole genome sequence of 134 *S*. Typhimurium U288 isolated from animals in years the 2014 and 2015 (APHA collection). To place these in the phylogenetic context of *S*. Typhimurium, we included 131 isolates that represented diverse phage types, including 12 isolates from the current monophasic *S*. Typhimurium ST34 epidemic (40). We also included the 33 human clinical isolates from 2014 and 2015 from the main U288 clade and 13 closely related isolates, and a U288 isolate (CP0003836), reported previously from Denmark in 2016 (41), for context. The phylogenetic structure of *S*. Typhimurium was consistent with that described previously (42), with a number of deeply rooted lineages, some of which exhibited evidence of clonal expansion at terminal branches (Figure 2). All *S*. Typhimurium U288 isolates from pigs were present in a single phylogenetic clade together with the 33 isolates from human clinical infections (U288 clade, green lineages, Figure 2). The U288 clade was closely related to thirteen *S*. Typhimurium isolates of various other phage types but none were phage type U288. Most of these were isolated from human clinical infections, and just two from animal hosts, both from avian hosts (Figure 2). Of note, *S*. Typhimurium strain ATCC700720 (LT2) differed by fewer than 5 SNPs from the common ancestor of the U288 clade and the 13 related non-U288 strains. *S*. Typhimurium strain ATCC700720 (LT2) was originally isolated from a human clinical infection at Stoke Mandeville hospital, London in 1948, and subsequently has been used for studying the genetics of *Salmonella* worldwide (43). The three U288 isolates from human clinical infections in the minor U288 clade clustered together with isolates of other non-U288 phage types.

**Figure 2.**
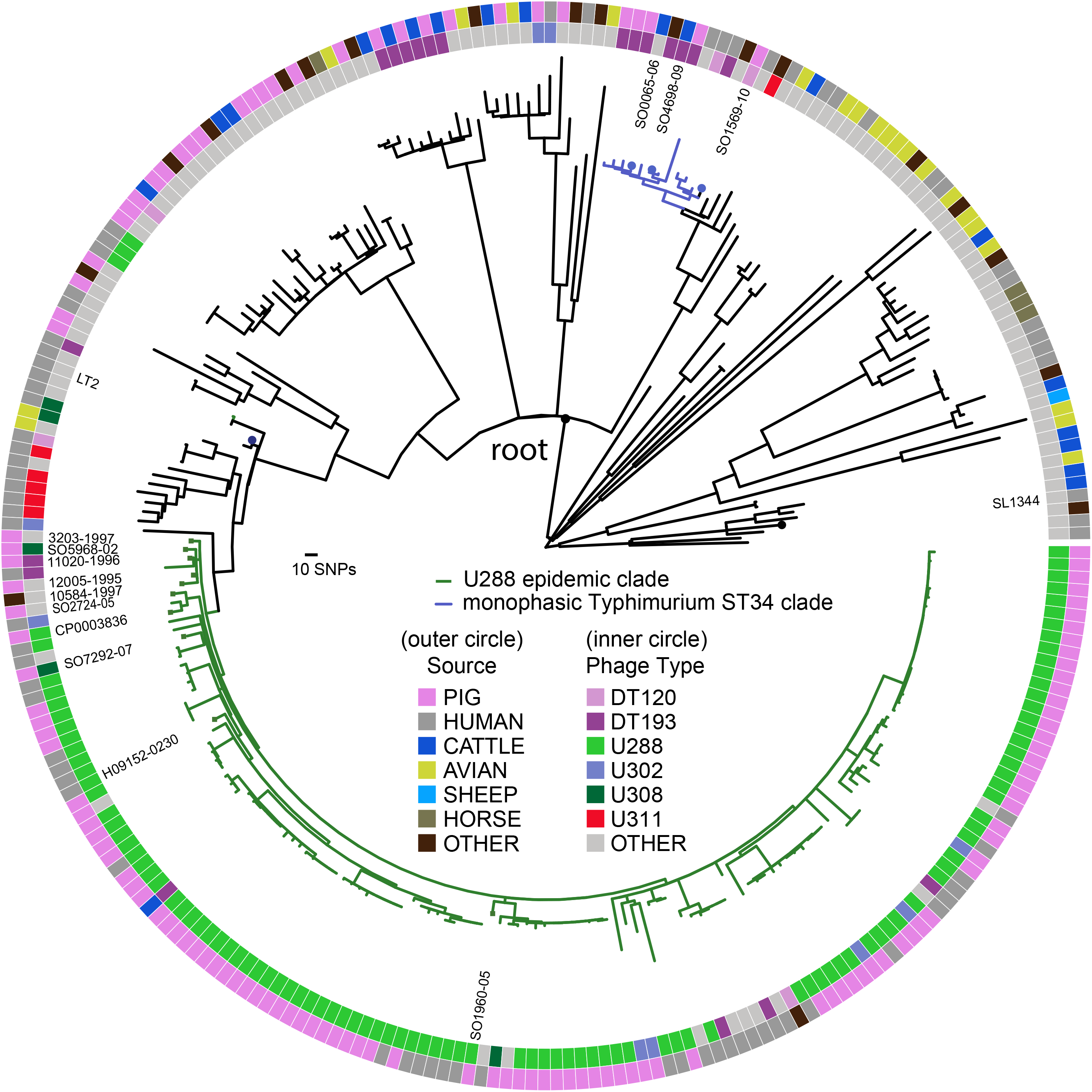
Phylogenetic relationship of *S*. Typhimurium U288 and *S*. 4,[5],12:i:- epidemic clades in the context of *S.* Typhimurium diversity. Maximum likelihood phylogenetic tree constructed using recombination-purged core genome sequence variation (7189 core SNPs) in the whole genome sequence of 265 *S*. Typhimurium isolates. The source of isolate (outer circle) and phage type of isolate (inner circle) are specified by the fill colour as indicated in the key. *S*. Typhimurium U288 (green lineages) and monophasic *S*. Typhimurium ST34 (blue lineages).

### *S*. Typhimurium U288 are rare and monophasic *S*. Typhimurium ST34 are common in human clinical cases

Epidemiological surveillance based on phage-typing indicated that *S*. Typhimurium U288 is rarely isolated from clinical cases of salmonellosis in the UK, while monophasic *S*. Typhimurium is commonly reported (7). A similar epidemiology has also been noted in other European countries, including Denmark (10). However, estimation of relative risk of human infection based on phage-typing is potentially misleading, due to potential polyphyletic clusters of common phage-types. Our phylogenetic analysis indicated that the majority of U288 isolates are from a single clonal group, but that this clade also contained a number of isolates that were not identifiable by phage typing as U288, therefore, the contribution to human infection may be underestimated. Conversely, a proportion of the *S*. Typhimurium U288 isolates from human clinical infections were only distantly related to the clonal group of *S*. Typhimurium U288 associated with pigs in the UK, and therefore, contribute to an over estimation of the pig-associated U288 genotype to human infection. We more accurately estimated the relative contribution of the U288 clade variant and monophasic *S*. Typhimurium ST34 variant to human infection by determining the relative contribution of the *S*. Typhimurium U288 clade isolates and monophasic *S*. Typhimurium ST34 clade isolates to human clinical infections in the UK between April 2014 and December 2015. Of 1666 *S*. Typhimurium isolated in this period 33 (1.9%) were from the U288 clade. In contrast, 894 isolates (54%) were from the monophasic *S*. Typhimurium ST34 clade.

### Distinct prophage repertoire, plasmid content and genome degradation of *S*. Typhimurium U288 strain S01960-05 and monophasic *S*. Typhimurium ST34 strain S04698-09

In order to compare the whole genome sequence of *S*. Typhimurium U288 and monophasic *S*. Typhimurium ST34, we assembled and closed the genome sequence of strain S01960-05, a representative U288 isolated in 2005 from a pig, using long-read sequence data. Alignment with monophasic *S*. Typhimurium ST34 strain S04698-09 indicated overall synteny, that was interrupted by seven insertions or deletions (indels) greater than 1 kb due to distinct prophage occupancy, recombination within shared prophage in one or other genome, the presence of a composite transposon encoding multidrug resistance, and an integrative conjugative element (SGI-4) in strain S04698-09 (Figure 3). Both genomes had Gifsy1, Gifsy2, ST64B, similar partial sequence of Fels-1 and remnant prophages SJ46 and BCepMu. The *thrW* locus was variably occupied by either the mTmV prophage in monophasic Typhimurium ST34 strain S04698-09 that carries the *sopE* gene (21), or ST104 in U288 strain S01960-05, a prophage previously described in *S*. Typhimurium DT104 strain NCTC13384 (44). U288 strain S01960-05 had a complete Fels-2 prophage which was absent from S04698-09. The S04698-09 genome also harboured two additional prophages related to HP1 and SJ46, that were absent from U288 strain S01960-05.

**Figure 3.**
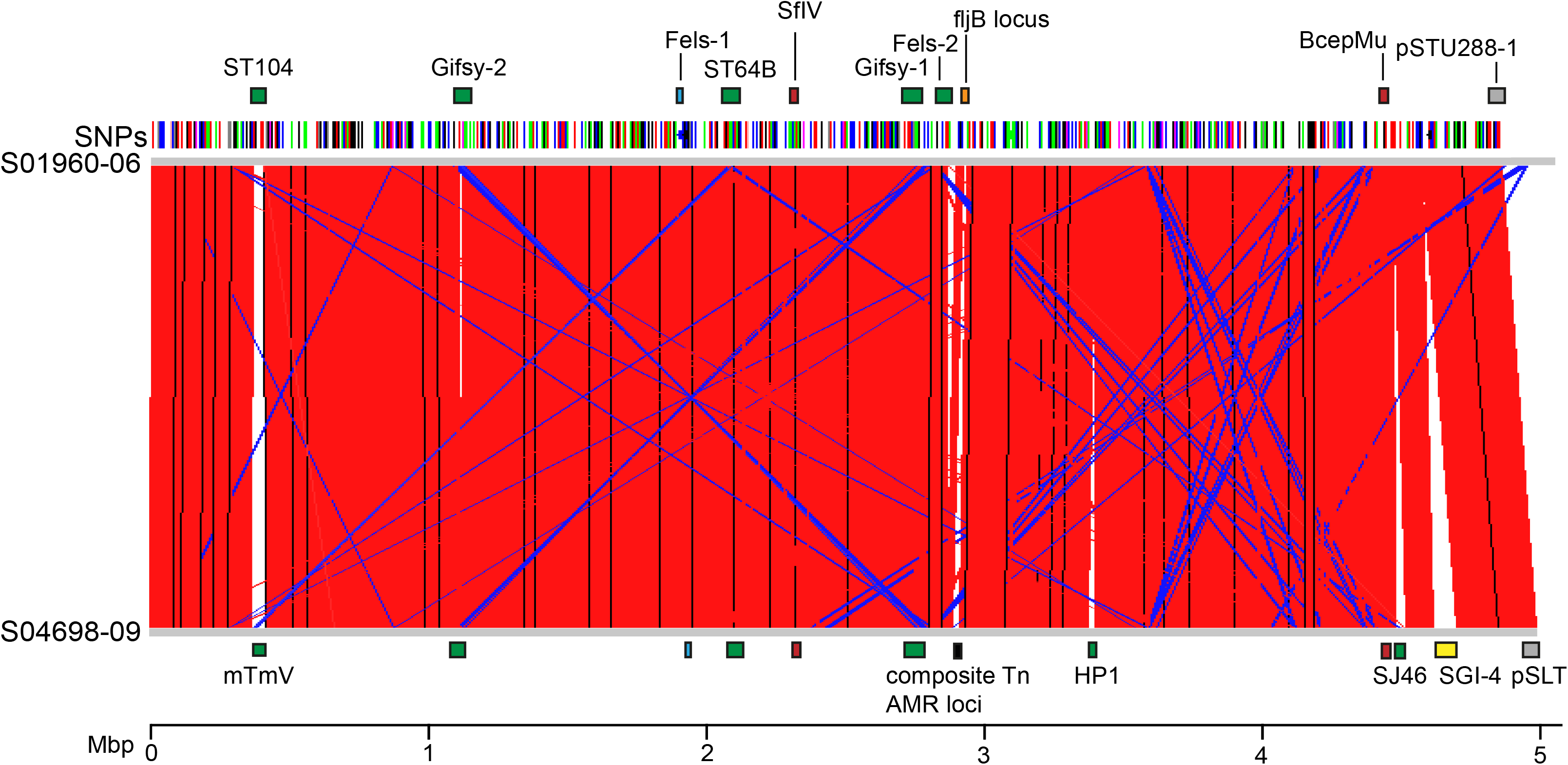
Comparison of *S*. Typhimurium U288 strain S01960-05 and *S*. 4,[5],12:i:- S04698-09 genomes. Red bars indicate sequence identity, blue bars indicate reverse and complement sequence identity >90%. Notable features highlighted by coloured bars and labels on respective genome (S01960 upper, S04698 lower). Features with no label are identical to their respective partner on the opposite genome. Prophage identified by PHAST software: green bars indicate predicted complete functional phage, blue bars indicate partial prophage, red bars indicate incomplete phage sequence. Purple bars denote plasmids. Yellow bar denotes genomic island, SGI-4. Orange bar represents the *fljB* locus, and the equivalent region in S04698 that is disrupted by composite transposon insertion encoding antimicrobial resistance (AMR) genes. Location of SNPs with reference to the S01960 fasta as a reference and S04698 fastq as a query sequence. Scale bar indicates the base location (Mbp).

Whole genome sequence analysis of a *S*. Typhimurium U288 strain previously identified three plasmids, pU288-1, pU288-2 and pU288-3 (41, 45). These three plasmids were also present in *S*. Typhimurium U288 strain S01960-05. In contrast, no plasmids were present in monophasic *S*. Typhimurium ST34 strain S04698-09. The pU288-1 is similar to the virulence plasmid pSLT, an IncF plasmid present in many *S*. Typhimurium strains (45). Additional sequence in pU288-1 but absent in pSLT was a composite transposon region containing integrons with AMR genes including *bla*_TEM_, *sulIII*, *aadA*, *cmlA*, *aadA2*, and *dfrA*. The IncQ1 plasmid pU288-2 encoded additional AMR genes *sulII*, *strA*, *strB*, *tetA* and *cat*.

A notable difference in coding capacity affecting the core genome of the two strains resulted from hypothetically disrupted coding sequences (HDCS) due to introduction of a premature nonsense codon from small insertions or deletions (indels) resulting in a frame shift or single nucleotide polymorphisms (SNP). The monophasic *S*. Typhimurium ST34 strain S04698-09 genome contained three HDCS outside of prophage, with reference to *S*. Typhimurium SL1344. In contrast, *S*. Typhimurium U288 strain S01960-05 contained 19 HDCS outside of prophage (Table 1). Nine U288 HDCS encoded hypothetical proteins of unknown function (*ygbE*, *yciW*, SL2283, *yfbK*, SL2330, *yhbE*, SL0337, *ybaO*, SL1627, *yfbB*, and *yqaA*), while ten had predicted functions based on sequence similarity or known functions (*assT5*, *assT3*, *dtpB*, *hutU*, *cutF*, *oadA*, *pncA*, *sadA*, *tsr*, *oatA*, and *rcoR*). Several of these may be important for colonisation of the caecum of pigs following oral inoculation based on an initial screen using transposon insertion library (46). In particular, several insertion mutants in *assT3* and *assT5* indicated a fitness score of −2 and −5.76.

**Table 1.** Hypothetically disrupted coding sequence (HDCS) in the S. Typhimurium U288 strain S01960-09 with reference to monophasic S. Typhimurium ST34 strain S04698-09. Locus tag is given, along with predicted function. The type of sequence polymorphism leading to truncation is indicated as insertion (ins), deletion (del) or non-sense codon (nsSNP) with reference to sequence (S04698-08). * Fitness score was derived from the screening of *S*. Typhimurium strain ST4/74 mutants by oral inoculation of pigs and recovery from the colon using transposon-directed insertion site sequencing (46). The fitness score is the log_2_-fold change in the number of sequence reads across the boundaries of a transposon insertion in the gene between the input and output pools, after normalisation to account for variations in the total number of reads obtained for each sample. Zero indicates no change in relative abundance, positive values indicate mutants that are more abundant in the output pool and negative values suggest attenuation. Where multiple mutants for a given gene were screened, the mean score is presented with the number of mutants shown.

| Polymer-phisms | SL1344 locus | Gene | Product | Fitness score* | No. of mutants |
| --- | --- | --- | --- | --- | --- |
| **del** | 0242 | *nlpE/cutF* | Copper homeostasis protein precursor/lipoprotein precursor | +1.26 | 2 |
| **ins** | 0337 | NA | Hypothetical periplasmic protein | -15 | 1 |
| **ins** | 0453 | *ybaO* | Hypothetical transcriptional regulator | - | 0 |
| **ins** | 0744 | *oadA2* | Oxaloacetate decarboxylase alpha chain | +0.41 | 3 |
| **ins** | 0767 | *hutU* | Urocanate hydratase | +1.19 | 1 |
| **ins** | 0987 | NA | Hypothetical host specificity protein (Gifsy-2) | +0.55 | 5 |
| **ins** | 1228 | *pncA* | Nicotinamidase/pyrazinamidase |  | 0 |
| **del** | 1627 | NA | Conserved hypothetical protein | -2.96 | 1 |
| **nsSNP** | 1632 | *yciW* | Conserved hypothetical protein | -0.87 | 1 |
| **ins** | 2208 | *oafA/oatA* | O-antigen acetylase/ O-acetyltransferase | -1.52 | 10 |
| **del** | 2274 | *menE* | O-succinylbenzoic acid-CoA ligase | -0.65 | 4 |
| **del** | 2282 | *elaC* | Ribonuclease Z | -4.03 | 1 |
| **del** | 2283 | *cheV* | Hypothetical receptor/regulator protein | - | 0 |
| **del** | 2284 | *yfbK* | Lipoprotein | +0.27 | 1 |
| **del** | 2285 | *nuoN* | NADH-quinone oxidoreductase subunit N | - | 0 |
| **ins** | 2330 | *rocR* | Hypothetical transcriptional regulator/Arginine utilization regulatory protein | +0.20 | 2 |
| **del** | 2661 | *bapA* | Beta-peptidyl aminopeptidase | +1.13 | 19 |
| **ins** | 2804 | *yqaA* | Putative membrane protein | +1.08 | 1 |
| **del** | 2911 | *ygbE* | Inner membrane protein | +2.43 | 1 |
| **ins** | 3165 | *assT3* | Probable arylsulfate sulfotransferase | -5.76 | 2 |
| **ins** | 3274 | *yhbE2* | Hypothetical membrane protein/ inner membrane transporter | -0.77 | 1 |
| **ins** | 3557 | *dtpB* | Dipeptide and tripeptide permease | - | 0 |
| **nsSNP** | 3656 | *sadA* | Autotransporter/adhesin | -0.08 | 8 |
| **del** | 3760 | *deoR2* | Transcriptional regulator | +0.66 | 1 |
| **nsSNP** | 3934 | *assT5* | Probable arylsulfate sulfotransferase | -2.05 | 4 |
| **ins** | 4464 | *tsr* | Methyl-accepting chemotaxis protein | - | 0 |

Analysis of the pangenome revealed a similar sized core and accessory genome, with 3962 and 4056 core gene families, out of pangenomes of 5501 and 5578 gene families, for U288 and ST34, respectively (Supplementary Figure 2). Lineage-specific differences in the accessory genome of U288 and ST34 were largely due to distinct prophage repertoires, the presence of SGI-4 in ST34, and the presence of pU288-1 (pSLT related) in U288. The prophage repertoire was particularly variable within the ST34 lineage, while plasmid sequence, including pU288-1 was highly variable in U288. Notably, U288 lacked lineage-specific genes present on the chromosome, with the exception of prophage.

### The *S*. Typhimurium U288 clade evolved from an *S*. Typhimurium LT2-like common ancestor by genome degradation and acquisition of AMR genes

To investigate the evolutionary events associated with the emergence of the *S*. Typhimurium U288 clade, using short read sequence data of U288 clade isolates and related *S*. Typhimurium isolates we determined the distribution of AMR genes, plasmid replicons and plasmid sequence (pSLT, pU288-1, pU288-2 and pU288-3), and identified allelic variants of genes identified as HDCS in *S*. Typhimurium U288 reference strain S01960-05 (Figure 4).

**Figure 4.**
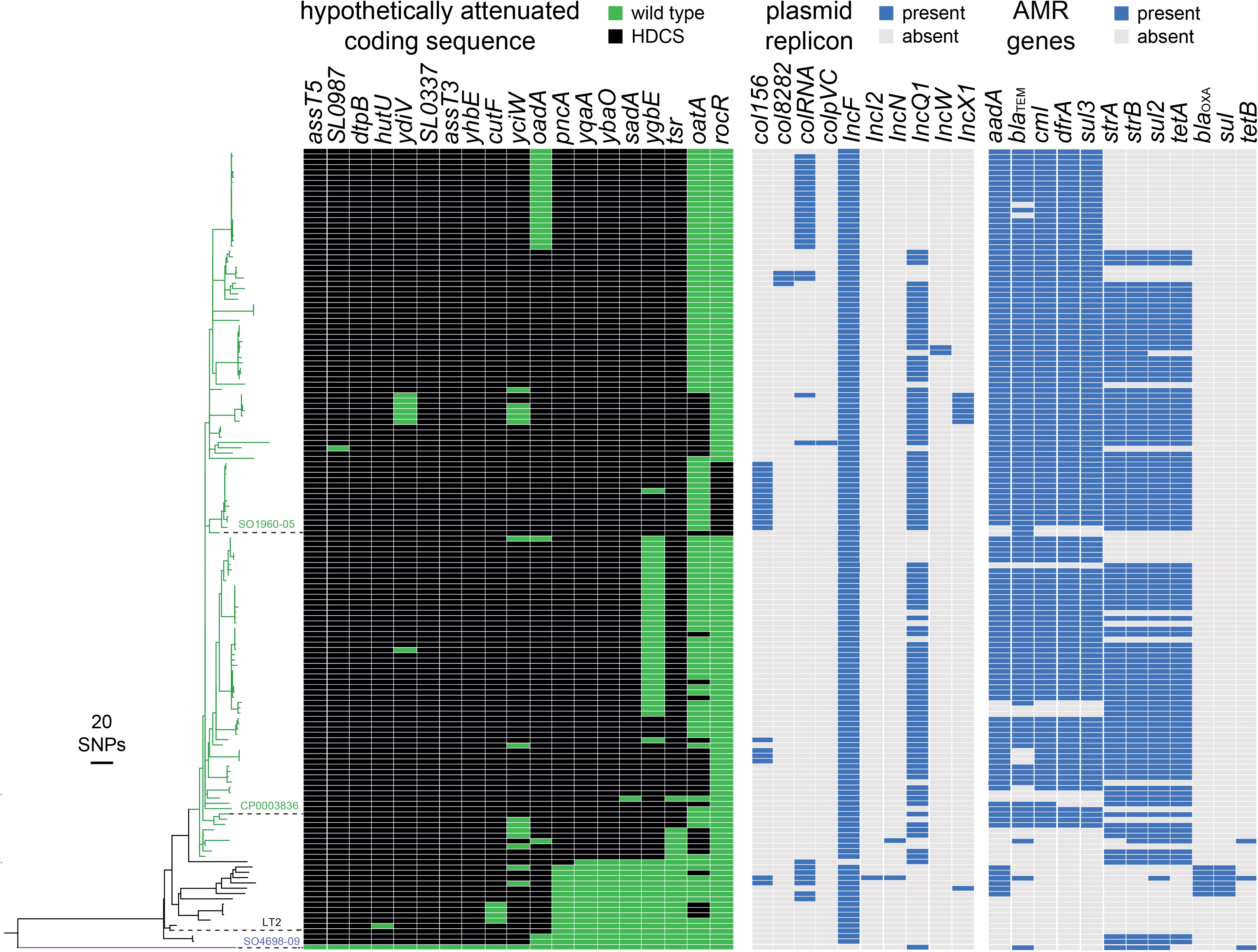
Phylogenetic relationship and genotypic variation in selected genes of *S*. Typhimurium U288 and monophasic *S*. Typhimurium ST34 strain S04698-09. Maximum likelihood tree based on recombination-purged variation in the core genome (1859 SNPs) with reference to *S*. Typhimurium strain LT2. ARIBA generated plasmid and resistance gene profiles are shown. Scale bar indicates the estimated number of SNPs based on the genetic distance.

The incF replicon was present in all but one isolate from the U288 clade and closely related isolates, consistent with the presence of all or part of the pSLT plasmid-associated sequence (Figure 4). Deletions of large parts of the pSLT-associated sequence were evident in the U288 clade isolates. Furthermore, in 58 of 133 U288 clade isolates, deletions affected two or more of the *spv* genes, previously implicated in virulence in the mouse model of infection (47).

The pattern of AMR gene presence was consistent with acquisition in two distinct evolutionary events. First, an incQ1 plasmid (pU288-2) was acquired concurrent with initial clonal expansion of the clade, followed by subsequent acquisition of a composite transposon on the pSLT-like plasmid, pU288-1 (Figure 4). The incQ1 replicon of the pU288-2 plasmid was present in 97 of 133 U288 clade isolates, including a cluster of six most deeply rooted isolates in the U288 clade, and was associated with the *strA, strB, sul2 and tetA* genes, encoding resistance to streptomycin, tetracycline and sulphonamide antibiotics. AMR genes *cml*, *sul3*, *dfrA, bla*_TEM_ and *aadA* that confer resistance to chloramphenicol, sulphonamides, trimethoprim, LJ-lactam and aminoglycoside antibiotics respectively were present on pU288-1 in all but 16 U288 clade isolates. These AMR genes were absent from six of the most deeply rooted isolates in the *S*. Typhimurium U288 clade.

Investigation of the distribution of the 19 HDCS identified in *S*. Typhimurium U288 strain S01960-05 within diverse *S*. Typhimurium strains in the context of their phylogenetic relationship indicated their sequential acquisition during the evolution of the U288 clade (Figure 4). Of the 19 HDCS, six (SL0337, *yhbE*, *cutF*, *yciW* and *oadA*) were also present in the genome sequence of closely related isolates including LT2, and five (*assT5*, SL0987, *dtpB*, *hutU* and *ydiV*) were present in isolates from two relatively distinctly related clades. Two additional HDCS in *S*. Typhimurium U288 strain S01960-05 and, were either sporadically present as HDRC throughout the *S*. Typhimurium collection (*oatA*), or only present as HDCS in strain S01960-05 and 14 closely related isolates. Six HDCS widely present in the U288 clade (*pncA*, *yqaA*, *ybaO*, *sadA*, *ygbE* and *tsr*), although a wild type allele of *ygbE* was present in a subclade of 35 isolates, and additional other isolates, suggesting that these may have subsequently reverted.

### AMR gene acquisition and genome degradation preceded the U288 epidemic clonal expansion

In order to investigate the temporal relationship between the emergence of the U288 epidemic in UK pigs around the year 2000 and the acquisition of AMR genes and genome degradation, we investigated the accumulation of SNPs on ancestral lineages and constructed a time scaled phylogenetic tree from variation in the core genome of U288 clade and genetically closely related isolates (Figure 5). To enhance the accuracy for the determination of the molecular clock rate, we supplemented the 131 strains from the year 2014 with 90 additional WGS of U288 isolated between 2006 and 2017. A maximum likelihood phylogenetic tree rooted with *S*. Typhimurium strain SL1344 outgroup was constructed from recombination-purged SNPs in the core genome. Root-to-tip accumulation of SNPs exhibited a molecular clock signal with a statistically significant fit to a linear regression model (R_2_=0.43, p<0.0001) (Figure 5A). A time dated tree was estimated in a Bayesian inference framework in order to determine the date of all nodes of the tree (Figure 5B). This analysis predicted the MRCA of all isolates at approximately the year 1937 (range 1915-1957), eleven years prior to the isolation of strain LT2 in London, and the MRCA of the U288 epidemic clade in 1988 (range 1982-1994). The disruption of the *pncA* gene was likely to have occurred between 1968 and 1979. Disruption of *ybaO*, *sadA*, *yqaA* and *ygbA* along with acquisition of pU288-2 between 1980 and 1990. Disruption of the *tsr* gene and the acquisition of the composite transposon that carried *cml*, *sul3*, *dfrA, bla*_TEM_ and *aadA* was acquired by plasmid pU288-1 was likely between 1993 and 1995. Disruption of the *rcoR* gene that only affected a subclade of U288 containing the reference strain S01960-05, likely occurred between 2000 and 2003.

**Figure 5.**
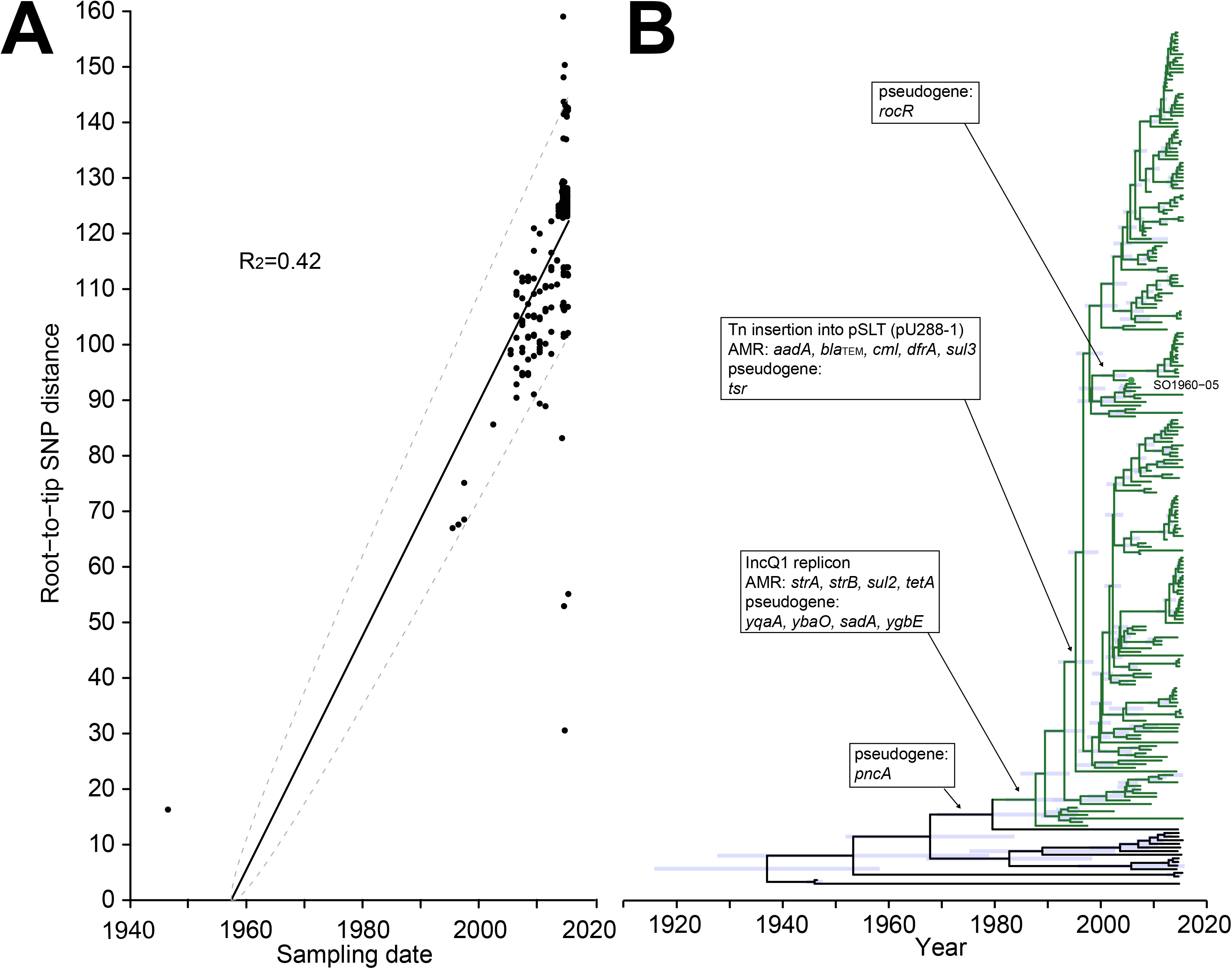
Time-scaled phylogenetic analysis of the emergence of the U288 epidemic clade. Population structure of 221 U288 clade strains isolated between 2005 and 2016, and 16 closely related strains isolated between 1947 and 2016 calculated using maximum likelihood estimation based on recombination-purged variation in the core genome sequence. (A) Linear regression of root-to-tip SNPs with a slope of 2.05 SNPs per year, and (B) estimated time-scaled phylogeny, with red bars indicating 95% CI of dated nodes, green lineages indicating the U288 epidemic clade, and black lineages indicating closely related lineages and clade. Major evolutionary events (arrows) of the acquisition of plasmids and transposons associated with AMR genes, and the accumulation of hypothetically disrupted coding sequences (HDCSs) resulting in possible pseudogene formation are indicated in boxes.

### *S*. Typhimurium U288 isolates have a longer doubling time and exhibit greater sensitivity to desiccation compared to ST34

Growth and survival in stress conditions encountered in food is likely to be an important factor in risk to food safety presented by *Salmonella* Typhimurium. We therefore compared U288 and ST34 isolate replication rate, motility, biofilm formation and ability to survive desiccation (Figure 6). Strains from the U288 clade exhibited a longer aerobic and anaerobic doubling time and increased sensitivity to desiccation, but similar motility and capacity to form biofilm, compared to monophasic *S*. Typhimurium ST34 isolates. The mean doubling time for in the U288 isolates was 0.6 hours and 0.54 hours in aerobic and anaerobic environments, respectively, compared to 0.52 hours and 0.47 hours for three ST34 isolates (Figure 6A).

**Figure 6.**
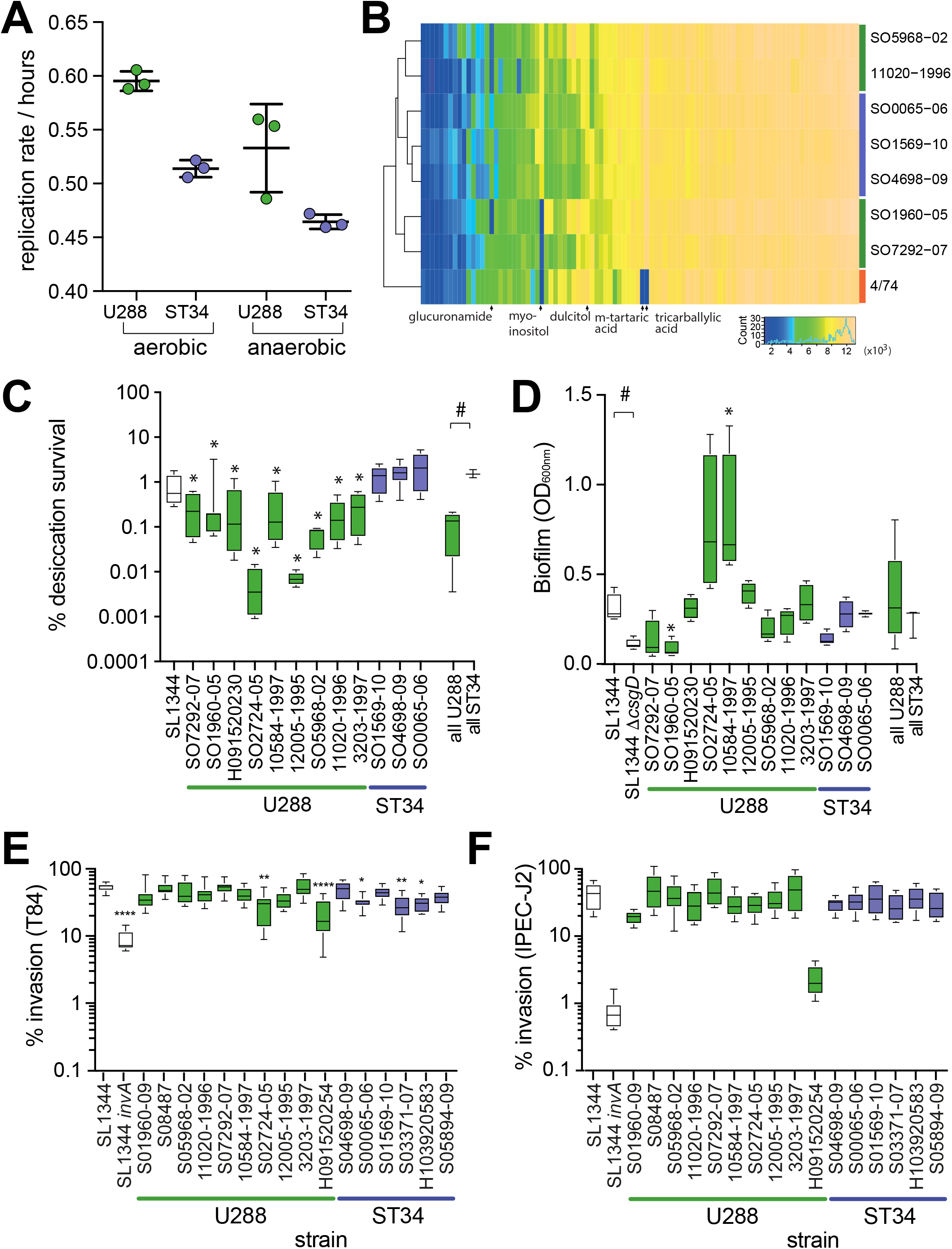
*In vitro* replication, carbon metabolism, sensitivity to desiccation and biofilm formation of *S*. Typhimurium U288 and ST34 isolates. (A) Circles indicate the mean doubling time of three S. Typhimurium U288 strains (S01960-09, S07292-07 and H09152-0230, green) and three ST34 strains (S04698-09, S00065-06 and S01569-10, blue) are indicated with the mean (horizontal bar) +/− with the standard error. (B) Metabolism measured using the BIOLOG phenotyping microarray platform in the presence of 95 carbon sources. The mean absorbance of at least two technical replicates were used to determine the area under the curve for each metabolite and are presented as a heat map with carbon sources in columns and eight S. Typhimurium strains, U288 (green bars), ST34 (blue bars) and strain 4/74 (orange bar) in rows. Unsupervised clustering of the metabolic activity of each strain and for each metabolite among test strains are indicated (left and above, respectively). (C) Proportion of CFU surviving desiccation for 24 hours with reference to the initial inoculum. The mean (horizontal line), interquartile (box), and range (vertical lines) are indicated. * Indicates mean survival that were significantly different from S04698-09, and # significantly different for strains compared as indicated by brackets, assessed using a Mann-Whitney U test of significance (p < 0.05). (D) Biofilm formation estimated from the measurement of biomass by crystal violet staining. Crystal violet retention measured by absorbance at 340nm with the mean (horizontal line), interquartile (box), and range (vertical lines) are indicated. * Indicates mean survival that was significantly different from S04698-09, and ^#^ indicates mean survival that was significantly different for strains compared indicated by brackets, assessed using a Mann-Whitney U test of significance (p < 0.05).

Since strains of each clade exhibited distinct replication rates, we compared respiration for three isolates of ST34 and four isolates of U288, utilizing a range of substrates as the sole carbon source for metabolism. All of the strains tested were able to use the majority of 95 carbon sources tested, but there was variation among approximately a quarter of substrates (Figure 6B). The pattern of carbon source utilisation of U288 and ST34 isolates was distinct from the commonly used lab strains *S*. Typhimurium 4/74 (histidine prototroph variant of strain SL1344). The inability or diminished ability of strain 4/74 to utilise m-Tartaric, Tricarballylic acid and D-xylose was a major factor that distinguished this strain from U288 and ST34 strains. Three ST34 strains clustered together, but the U288 isolates exhibited considerably greater diversity in carbon source utilization. Utilization of myo-inositol as a sole carbon source was the most pronounced phenotype that distinguished the two clusters of U288 isolates. Of note, strain S05968-02 and 11020-1996 that were able to use myo-inositol were isolated earlier and were more deeply rooted than strains S01960-09 and S07292-07 that were unable to use this source of carbon.

Desiccation is a common stress associated with the food chain. We observed a clade-specific variation in tolerance to desiccation by comparing ten U288 strains and three ST34 strains (Figure 6C). Following desiccation for 24 hours, approximately 2% of the initial inoculum remained viable for all three ST34 strains. In comparison the mean viability of U288 strains was 0.1%, but varied between 0.0001% and 0.3% among the ten U288 strains tested.

The loss of ability to form biofilm is a common feature of some host-adapted variants of *Salmonella enterica* (48, 49) (Figure 6D). *S*. Typhimurium strain SL1344 formed moderate biofilm that was dependent on expression of the *csgD* gene as previously described (48). The mean biofilm formation for ten U288 strains was not significantly different from that of three ST34 strains. However, considerable variation was observed especially for the U288 strains, and two strains of U288 had a statistically significant difference in biofilm formation compared to ST34 strain S04698-09, with U288 strain S01960-05 produced significantly less biomass and strain 10584-1997 significantly greater biomass.

### *S*. Typhimurium U288 and monophasic *S*. Typhimurium ST34 isolates exhibit distinct interactions with the host

We initially evaluated the interaction of representative strains of ST34 and U288 with tissue culture cells. No difference in the ability of U288 or ST34 isolates to invade human (T84) or porcine (IPEC-J2) epithelial cells in culture was observed (Figure 6E and 6F). Although, several strains of U288 and ST34 had a small but significantly lower invasion compared to strain SL1344 in T84 cells, and two U288 isolates (S01960-09 and H091520254) exhibited decreased invasion of IPEC-J2 cells.

In an initial experiment to investigate the host-pathogen interactions of U288 and ST34 isolates, we used the streptomycin pre-treated C57bl/6 mouse that is commonly used as model of *Salmonella*-induced colitis. We determined viable counts, pathology in the cecum and the expression of the Cxcl1 gene encoding the chemokine KC, and the Nos-2 gene encoding the innate immune effector iNOS, in cecum tissue three days after oral gavage with a single inoculum of one of five isolates of either U288 or ST34 (Supplementary Figure 4). Mice infected with strains of the same phylogroup (ST34 or U288) exhibited similar colonisation level (Supplementary Figure 3A), induction of KC and iNOS (Supplementary Figure 3B) and intestinal pathology (Supplementary Figure 3C). However, in each case, U288 strains colonised the mouse cecum to a greater level, induced higher levels of KC and iNOS transcripts and triggered a more severe pathology, compared to that resulting from infection with ST34 strains.

To compare the ability of six strains to colonise pigs, three U288 and three ST34 were modified by insertion of unique sequence tags in the chromosome to facilitate identification by sequencing, and four pigs were inoculated orally with an equal mixture of all six isolates. Colonisation was investigated after 48 hours by sequencing cultured homogenates of faeces and tissue and enumeration of the sequence reads for the strain-specific tags (Figure 7A). U288 and ST34 exhibited a distinct pattern of colonisation. ST34 isolates were more abundant than U288 isolates in three of four faeces samples on day 24 and 48 hours post inoculation. Isolates of neither variant were consistently dominant in the distal Ileum. In contrast, the U288 isolates were more abundant in the mesenteric lymph nodes and the tissue of the spiral colon.

**Figure 7.**
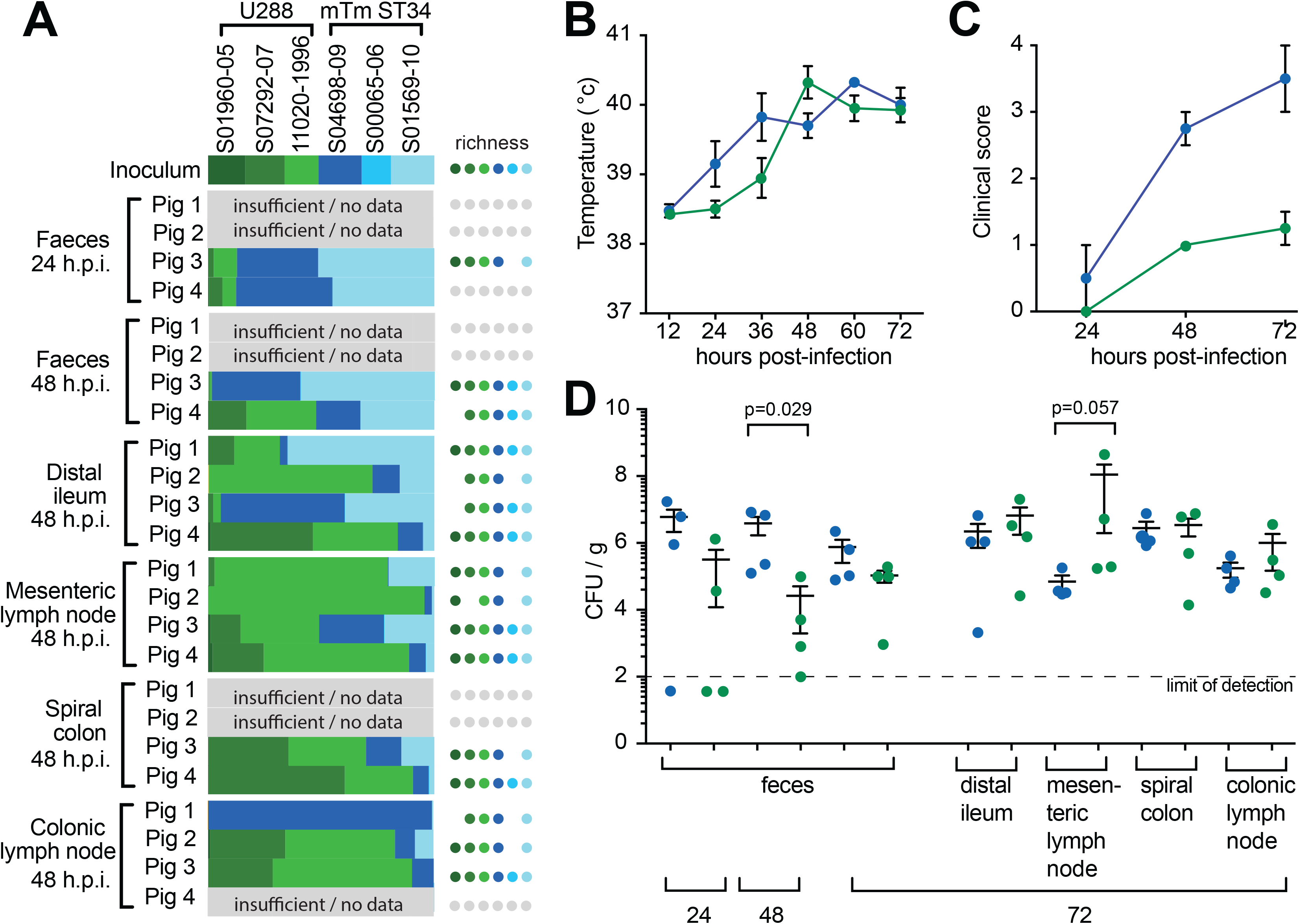
Colonisation and clinical signs of disease following oral inoculation of pigs with S. Typhimurium U288 and ST34. Four pigs were challenged orally with an inoculum containing six wild-type independently tagged strains (WITS) of *S*. Typhimurium (three DT193 and three U288) in approximately equal proportions (A). Whole genome sequence of the population of the WITS in the inoculum and recovered from infected pigs was used to enumerate each strain based on the unique sequence tag. The relative abundance of each strain is denoted by bars and the number of strains in each sample (richness) is indicated by circles based on the colour code indicated. Separately, four pigs were challenged orally with a strain of either ST34 strain S04698-09 or 11020-1996 (U288) to investigate the signs of disease and colonisation (B, C and D). The inocula used to challenge the pigs contained approximately 1×10^10^ colony forming units of either S04698-09 (DT193) or 11020-1996 (U288). Rectal temperatures of the pigs were monitored over 72 hours of infection (B). Clinical scores derived from physiological signs and faecal consistency (C). Viable counts of bacteria shed in the faeces and 24, 48 and 72 hours post inoculation and tissues collected 72 hours post-inoculation.

To further investigate the colonisation of pigs after oral inoculation, groups of four pigs were inoculated with either U288 strain S011020-1996 or ST34 strain S04698-09 in single infection experiments (Figure 7B to D). Rectal temperatures and clinical scores were recorded for all pigs throughout the infection, and at 72 hours post-inoculation the pigs were sacrificed, and colonisation of the faeces and tissues determined. Pigs inoculated with the U288 strain had significantly lower body temperatures at early time points post inoculation (Figure 7B), and this was reflected in significantly lower clinical scores than those inoculated with the ST34 strain (Figure 7C). The mean *Salmonella* CFU present in faeces was also lower at all time points in pigs inoculated with U288, compared with ST34 (Figure 7D). At 48 hours post-inoculation approximately 100-fold more ST34 than U288 were present in the faeces. Similar colonisation for each isolate was observed in the distal ileum and spiral colon, but U288 was present in higher numbers in the mesenteric and colonic lymph nodes (Figure 7D). These data were consistent with the mixed inoculum experimental infections.

## Discussion

The majority of *S*. Typhimurium U288 isolates from livestock and human infections were present in a phylogenetic clade that evolved from a common ancestor that was closely related to strain LT2. LT2 was isolated at Stoke Mandeville hospital in 1948 and has been used in a large number of studies on the genetics and biochemistry of *S.* Typhimurium (50). Approximately 50 years elapsed since the isolation of LT2 before U288 was first detected by epidemiological surveillance in the UK pig population, around the year 2003 (3). The zoonotic source of the infection caused by the LT2 strain and that of the common ancestor with U288, is not known (50). Evolution of successive descendants of the LT2-like hypothetical ancestor gave rise to multiple lineages evident from our analysis, including one that gave rise to the U288 epidemic clade in pigs. In our strain collection, non-U288 clade isolates that were also direct descendants of the LT2-like hypothetical ancestor, were isolated from human infections or from avian hosts (red lineages in Figure 2). Comparative genome analysis revealed that the evolution of U288 was characterized by the step-wise the acquisition of genes involved in resistance to multiple antibiotics, and the accumulation of genome sequence polymorphisms, some of which resulted in interruption of coding sequences.

The acquisition of resistance to antimicrobials is a key factor in the emergence of bacterial pathogens over the past 50 years, and is often the evolutionary event immediately preceding the spread and clonal expansion of a new clone (22, 51). A U288 isolate was previously reported to encode antimicrobial resistance genes on two plasmids, a pSLT-like plasmid pU288-1 with *tetA*, *sulII*, *strA* and *strB* genes, and an incQ plasmid pU288-2 (45). Resistance genes on pU288-1 resulted from acquisition of two mobile genetic elements, *bla*_TEM_ associated with an IS26 element, and *sulIII*, *aadA*, *cmlA* and *dfrA* associated with a class I integron in a composite Tn21-like element (45). The pU288-2 plasmid may have been acquired first as it is present in isolates from throughout the U288 clade. Insertions carrying AMR genes in pU288-1 may have occurred later than the acquisition of pU288-2, since a group of seven U288 isolates that form a more deeply rooted, basal clade, lacked the pU288-1-associated AMR genes. The majority of the U288 isolates in our analysis were direct descendants of the hypothetical ancestor that acquired AMR genes on pU288-1, and few from that of the basal clade that lacked these genes, suggesting pU288-1 evolution was an important event in the success of the U288 epidemic clade. However, despite a number of examples of apparent loss of AMR genes associated with both pU288-1 or loss of the pU288-2 plasmid (loss of AMR genes and the incQ replicon), there were just three isolates that had lost both concurrently, highlighting the importance of MDR.

Sequence polymorphisms in *S*. Typhimurium U288 strain S01960-05 resulted in truncated coding sequences affecting 26 genes, with reference to strain SL1344, a commonly used lab strain. Sixteen of these polymorphisms were predicted to have occurred before the hypothetical LT2-like ancestor of the U288 and related clades (red and green lineages in Figure 2). Six coding sequences (SL0337, *assT3*, *yhbE*, *cutF*, *yciB* and *yciW*) were truncated in all descendants of the hypothetical LT2-like ancestor, and one other (*oadA*) in all but one deeply rooted lineage. However, eight genes (*pncA*, *yqaA*, *ybaA*, *sadA*, *ygbE*, *tsr*, *oatA* and *rcoR*) were truncated in either the hypothetical ancestor of all U288 clade isolates, or subsequently in a subset of descendant lineages, in a stepwise manner. The first gene to be truncated was *pncA*, an event on an internal lineage on the phylogenetic tree that coincided with the acquisition of pU288-2, encoding AMR genes. Truncation of *pncA* is therefore, characteristic of the U288 clade. PncA is a nicotinamidase, a component of one of the pyridine nucleotide cycle (PNC) pathways, involved in the recycling nicotinamide adenine dinucleotide (NAD). The PncA dependent PNC pathway is probably more active in scavenging of pyridine compounds present in the environment (52). NAD is central to metabolism in all living systems, participating in over 300 enzymatic oxidation-reduction reactions (52), and therefore, conditions where de novo synthesis of NAD is limited by the availability of tryptophan or aspartate, the inability to use exogenous pyridines may limit metabolism. Perhaps significantly, the *pncA* gene is also truncated in *S*. Choleraesuis, a serotype that is highly host-adapted to pigs and replicates more slowly than *S*. Typhimurium in the pig intestinal mucosa (53). Chronologically, the next events were the truncation of the *yqaA*, *ybaA*, *sadA* and *ygbE* genes. SadA is a surface localised adhesin that contributes to cell-cell interactions and therefore multicellular behaviour (54). Truncation of *tsr*, that encodes a methyl accepting chemotaxis protein involved in energy taxis and colonisation of Peyer’s patches in the murine model of infection, was acquired by a common ancestor of the majority of the U288 clade. This polymorphism occurred on an internal branch of the tree that coincided with the acquisition AMR genes inserted on pU288-1 and a clone that spread successfully through the pig population.

Accumulation of SNPs by isolates since the hypothetical LT2-like common ancestor was approximately linear facilitating estimation of dates for nodes within the phylogenetic tree. This indicated that the truncation of the *pncA* gene, the earliest identified evolutionary event since the hypothetical LT2-like common ancestor specific to the U288 clade likely occurred in the 1970s, and culminated with the acquisition of AMR genes on pU288-1 and truncation of the *tsr* gene around 1995. The U288 clade may therefore have been evolving in the pig population before its first detection and rapid spread around the year 2003. The slow emergence of the U288 clade may have been associated with a gradual adaptation to a unique niche in the pig population, but it may also reflect a lag in time from emergence to detection by surveillance, as was proposed for other epidemics such as *S*. Enteritidis in poultry layer flocks, after the eradication of *S*. Gallinarum (55).

*S*. Typhimurium U288 and monophasic *S*. Typhimurium ST34 exhibited important differences in the way it interacted with the non-host environment, that could affect the likelihood that it survives in food and is transmitted on to consumers. Considerably more viable monophasic *S*. Typhimurium ST34 bacteria were recovered following desiccation for 24 hours, compared to U288. Many foodborne disease outbreaks due to *Salmonella* have been traced back to low moisture, ready-to-eat (RTE) foods (56), including dried pork products (57). Resistance to desiccation may be of particular significance because low oxygen tension is associated with increased resistance to a number of secondary stressors such as pH, salt, alcohol and heat (56). Monophasic *S*. Typhimurium ST34 also replicated at a significantly higher rate than U288 in culture, a characteristic that may result in a greater risk were it to apply in relevant food matrices. The infective dose of *Salmonella enterica* required to produce disease symptoms in humans? is in the range of 10^5^ to 10^9^ CFU, while naturally contaminated meat samples typically contained around 10^3^ CFU/g (58), suggesting that replication in food may be important for transmission. Enterobacteriaceae replicate in food stored at 7°C, increasing up to 10,000-fold in sausage meat in 2 weeks (59).

*S*. Typhimurium U288 and monophasic *S*. Typhimurium ST34 are associated with distinct risk to human health, despite circulating in the same pig population in the UK (3). Our data suggest that this may be due to the differences in tissue tropism and levels of each variant in the pig host that has the potential to impact the likelihood that *Salmonella* enters the food during the slaughter and butchering process. The greatest risk to *Salmonella* entering pork products is the contamination of the carcass at slaughter due to careless evisceration processes and inadequate cleaning of polishing machines (60). The relative contribution of *Salmonella* present in the faeces and tissues to contamination of food is not known, but increased colonisation of *Salmonella* in the caecum due to longer time in lairage correlated with contamination of the carcass at slaughter (61). Our data is consistent with a greater role for faecal contamination, since the mean viable counts of U288 isolates were up to two orders of magnitude lower in the faeces, but 10-fold greater in mesenteric lymph nodes, compared to monophasic *S*. Typhimurium ST34. The reason for the difference in colonisation of pigs is not known. However, monophasic *S*. Typhimurium ST34 lack the pSLT virulence plasmid, encoding the *spv* locus that is required for invasive disease in mice and severe gastroenteritis in cattle (62–64). Polymorphisms in some U288 strains also affect this locus. Acquisition of the IS26 element appears to have been accompanied by deletion of the *spvR* and *spvA* genes of pSLT, likely affecting expression of *spvB* and *spvC* that are pU288-1 also contained large regions of duplicated sequence of pSLT origin and sequence similar to other plasmids previously reported in *Salmonella* and *E. coli*, suggesting a complex history of horizontal gene transfer and deletion. The pU288-2 plasmid in U288 isolates acquired resistance genes in two distinct events, on an incQ plasmid pU288-2, and by acquisition of genes located on the pSLT-like plasmid pU288-1.

Taken together, our data contribute to a better understanding of the evolutionary history and phenotypes associated with the emergence of a new bacterial pathovariant. In the case of *S*. Typhimurium U288, emergence in pigs appears to have been associated with a decreased risk to food safety for the consumption of pork or cross-contaminated food products by the human population. However, the consequences to the health and productivity of pigs as a result of a more invasive disease is not known, but may be an important consideration for the pork production industry.

## Acknowledgements

Ethical approval for the experimental mouse infections was granted following review by the Animal Welfare and Ethical Review Body (AWERB, University of East Anglia, Norwich, UK) under project license PIL 70/8597. Ethical approval for experimental infections of pigs was granted following review by the Moredun Research Institute Ethical Review Committee under project licence PCD70CB48. All data is freely available in publicly accessible data bases under accession numbers reported in Supplementary Information. None of the authors have any competing interests to declare. RK was supported by research grants BB/N007964/1 and BB/M025489/1, and by the BBSRC Institute Strategic Programme Microbes in the Food Chain BB/R012504/1 and its constituent projects BBS/E/F/000PR10348 and BBS/E/F/000PR10349. MS was supported by BBSRC grant BB/M021114/1 and the Institute Strategic Programme Control of Infectious Diseases (BBS/E/D/20002173). NH was supported by BBSRC grant BB/M025411/1. This research was supported in part by the Norwich Biosciences Institute (NBI) Computing infrastructure for Science (CiS) group.

## Supplementary figures

**Supplementary Figure 1. Maximum likelihood estimation of the phylogeny of 1697 S. Typhimurium and monophasic S. Typhimurium isolated from human clinical infections in England and Wales in 2014-2015**.

**Supplementary Figure 2. Analysis of the pangenome of S. Typhimurium U288 and ST34**.

**Supplementary Figure 3. Invasion of epithelial cells in culture**.

**Supplementary Figure 4. Colonisation of C57bl/6 mice and immune response following oral inoculation with *S*. Typhimurium U288 or ST34**.

