## Supplementary figures and images for "Ecological niche adaptation of a bacterial pathogen associated with reduced zoonotic potential"

### Supplementary Figure 1

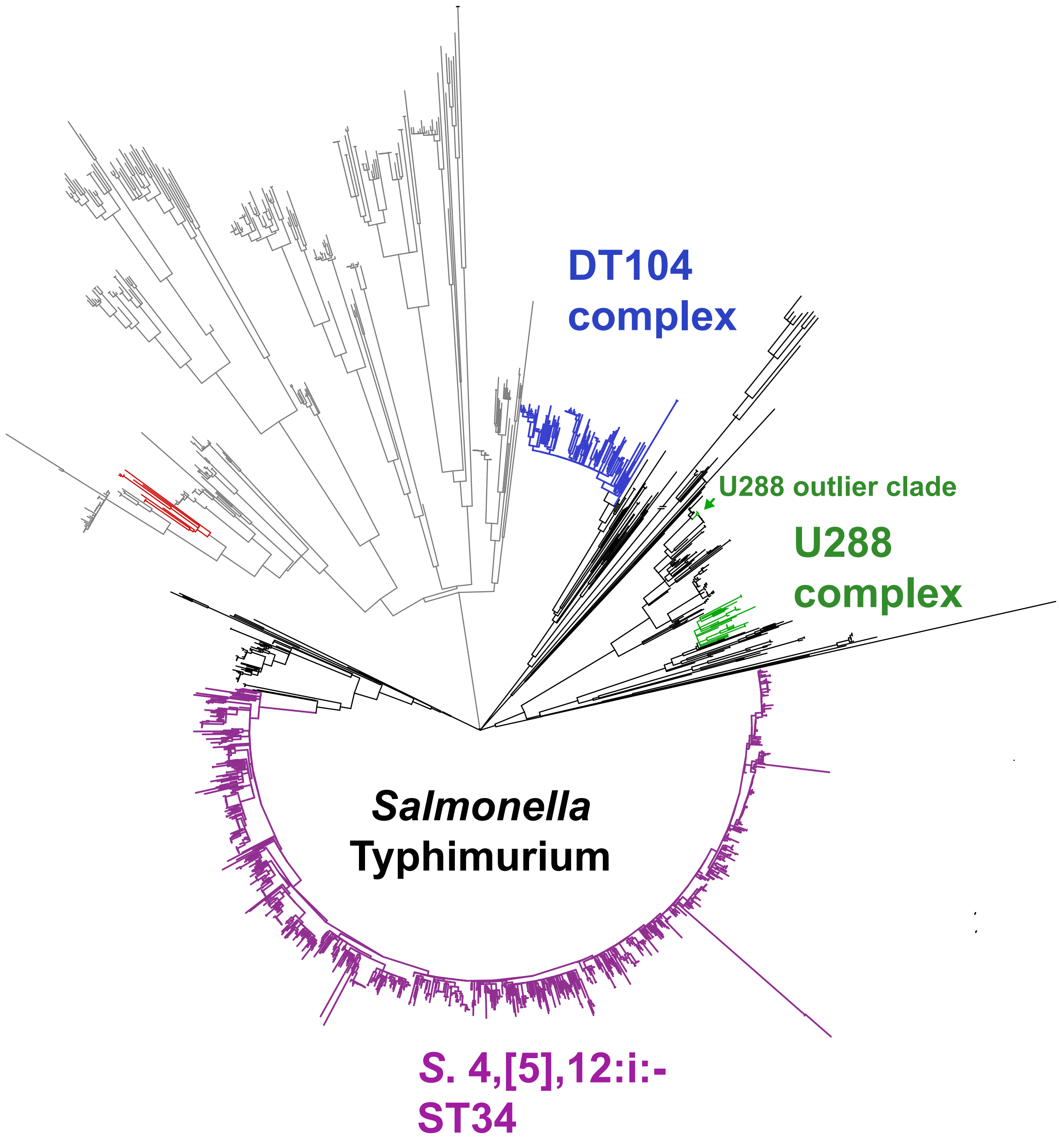

### Supplementary Figure 2

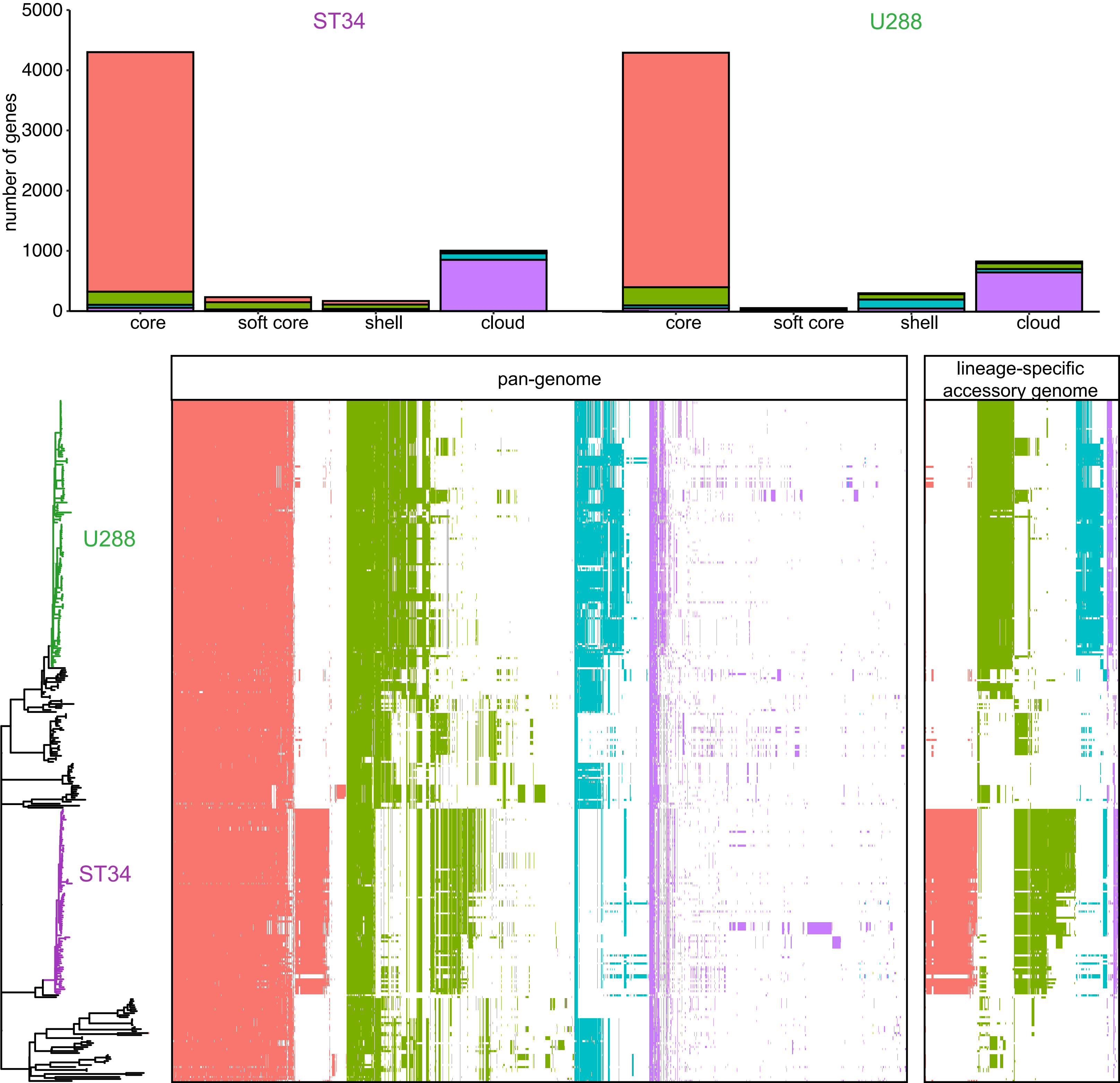

### Supplementary Figure 3

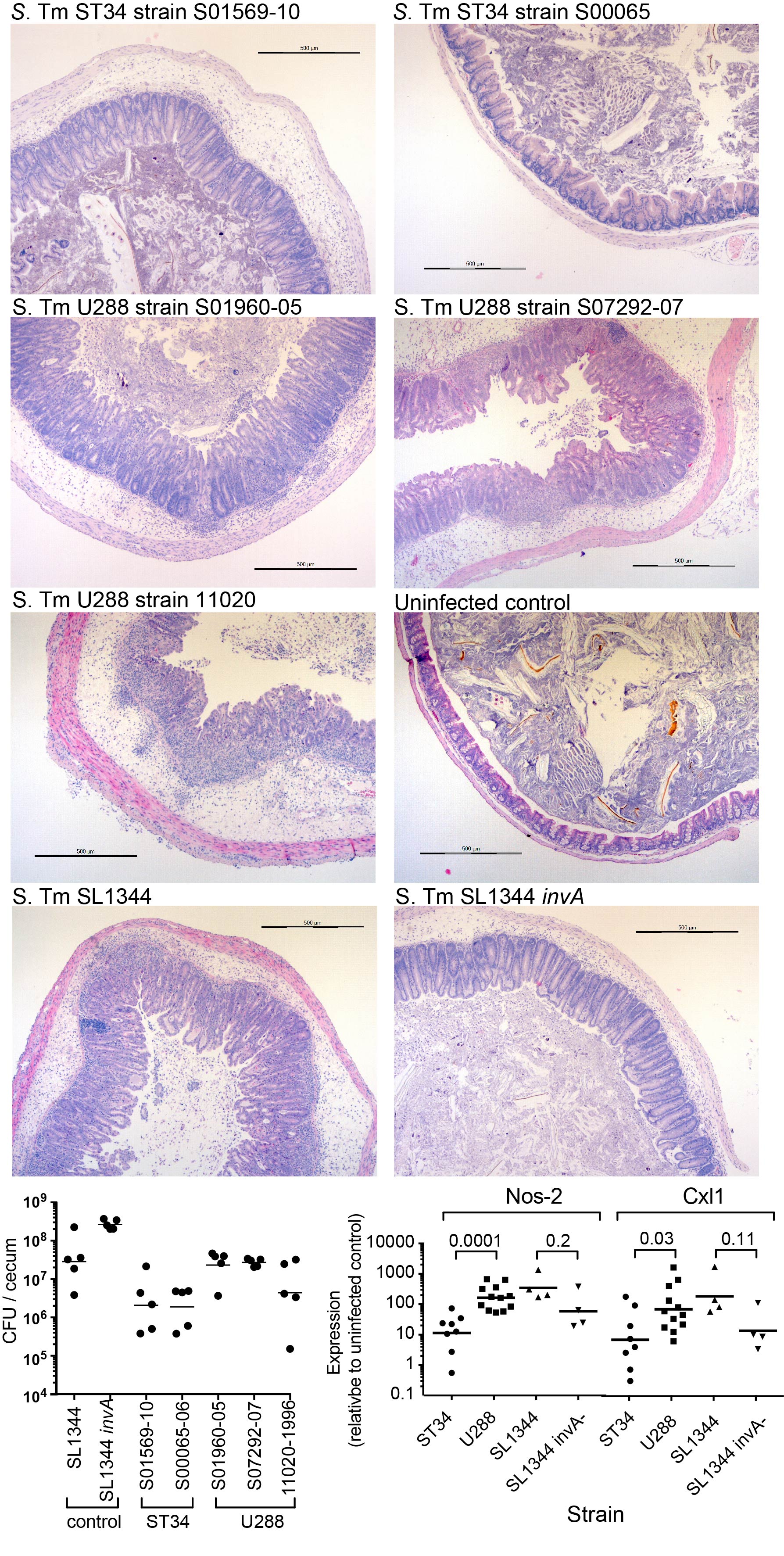
